# Surface charge tuning of lipid–polymer hybrid nanoparticles for optimized cilostazol delivery and platelet compatibility

**DOI:** 10.64898/2026.09.23.753835

**Authors:** María Francisca Matus, Bruno A. Cisterna, Marcelo M. Mariscal, Claudia Carranza, Martín Ludueña, S. Alexis Paz, Javier Salazar-Muñoz, A. Said Álamos-Musre, Eduardo Fuentes, Joel B. Alderete, Esteban F. Durán-Lara, Ricardo I. Castro, Iván Palomo, Cristian Vilos

**Author notes:** Corresponding authors. E-mail addresses (B.A. Cisterna), (I. Palomo), (C. Vilos).

## Abstract

Cardiovascular diseases (CVD) are the leading cause of morbidity and mortality worldwide. This clinical burden is predominantly attributable to atherothrombotic events resulting from platelet hyperactivation, a process integral to the pathophysiology of atherosclerosis. Cilostazol (CLZ) is a clinically established phosphodiesterase-IIIA inhibitor with antiplatelet and vasodilatory activity. However, its extremely low aqueous solubility results in dissolution-limited and variable oral absorption, while dose-dependent adverse effects and cardiovascular safety restrictions may constrain its clinical use. Lipid–polymer hybrid nanoparticles (LPHNPs) represent a highly promising drug delivery system to address these biopharmaceutical limitations. However, their interactions with blood components remain poorly understood, with surface charge among the most controversial physicochemical parameters implicated in platelet toxicity. Here, we applied an integrated *ex vivo* and *in silico* screening strategy to design LPHNPs based on poly(lactic acid) and poly(ethylene glycol)-modified lipids displaying positive (amine), nominally neutral (methoxy), and negative (carboxyl) surface charges to systematically identify safe boundaries at the nano–human platelet interface and evaluate their suitability as CLZ carriers. Positively charged nanoparticles precipitated immediately during self-assembly due to uncontrolled ionic pairing and were therefore structurally unviable. Neutral nanoparticles maintained a stable size (∼113 nm hydrodynamic diameter) but unexpectedly induced spontaneous platelet aggregation, limiting their applicability. In contrast, negatively charged nanoparticles exhibited exceptional colloidal stability (hydrodynamic size of ∼120 nm and ζ-potential of −48 mV), achieved significant inhibition of ADP-induced platelet aggregation, and adhered to the platelet surface without inducing activation or membrane disruption. Steered molecular dynamics simulations, performed under identical non-equilibrium pulling conditions, showed the highest interfacial resistance to CLZ extraction for the negatively charged model, qualitatively consistent with the sustained CLZ release profile observed *in vitro*. These findings highlight the importance of precise surface charge tuning in the development of safe cardiovascular nanomedicines based on LPHNPs, emphasizing the need for comprehensive platelet compatibility testing before further biological evaluation.

## 1. INTRODUCTION

Cardiovascular diseases (CVD) remain the leading cause of morbidity and mortality in the world [1,2], and their pathophysiology is closely related to atherosclerosis, a chronic inflammatory disorder of the arterial wall [3,4]. Platelets function as critical mediators in this landscape, participating not only in physiological hemostasis but also exacerbating pathology by driving acute atherothrombotic occlusions following atherosclerotic plaque rupture [5–8]. When endothelial damage occurs, platelets adhere to the site of injury, and the platelet plug formation occurs to prevent excessive bleeding. This process is triggered by a highly regulated set of physiological responses, including platelet adhesion, activation, secretion, and aggregation [9]. This cascade triggers multiple intracellular signaling pathways that induce platelet structural rearrangement and facilitate further recruitment [10]. Consequently, when this response occurs aberrantly at the site of atherosclerotic plaque rupture, it becomes the primary driver of atherothrombotic disease.

Antiplatelet drugs are widely prescribed for primary and secondary prevention of thrombotic events [11,12]. Cilostazol (CLZ), a quinolinone derivative, is a platelet anti-aggregant and vasodilator used for the treatment of intermittent claudication and for the secondary prevention of cerebral infarction [13–15]. It acts as an inhibitor of phosphodiesterase-IIIA (PDE3A) [16], leading to elevation of intracellular cyclic adenosine monophosphate (cAMP) levels in platelets and blood vessels [17,18]. Although CLZ is highly effective, the major drawbacks of its clinically available oral dosage form are the low water solubility (<6 μg/mL), poor absorption, and low oral bioavailability [19]. This limitation leads to requiring high daily oral doses that frequently provoke severe side effects [20].

The use of nanoparticles as drug delivery systems has emerged as a potent strategy to improve the bioavailability of poorly soluble drugs [21,22]. To date, diverse nanoparticles have been explored to increase the bioavailability of CLZ, including nanoemulsions [23,24], solid lipid nanoparticles [25], and polymeric nanoparticles [26]. Among these varied platforms, lipid– polymer hybrid nanoparticles (LPHNPs) have become the next-generation nanocarriers since they synergistically combine the high stability and controlled release kinetics of a rigid polymeric core with the biomimetic surface features and outstanding pharmacokinetic properties of lipid components [27–30]. These features make LPHNPs particularly attractive for the delivery of extreme hydrophobic cardiovascular agents such as CLZ, for which controlled release, improved bioavailability, and prolonged circulation are desirable therapeutic attributes. However, despite the favorable biocompatibility generally attributed to engineered nanocarriers, several studies have demonstrated that their entry into the vascular compartment triggers complex nano–blood cell interactions [31]. Notably, the ability of nanoscale physicochemical parameters, including particle size, surface chemistry, and surface charge, to unexpectedly alter platelet function is a systemic phenomenon inherent to all nanomaterials, regardless of their core composition or structural class [32–34]. Indeed, platelet responses to nanoparticles remain highly controversial, with studies across multiple nanomaterials describing both pro-aggregatory and apparently inert effects [35–37]. Surface charge is one of the most debated determinants of these outcomes. Positively charged nanoparticles often exhibit strong electrostatic interactions with platelet membranes, whereas neutral and negatively charged nanoparticles have produced inconsistent observations, ranging from mild activation to significant platelet aggregation [36–38]. These discrepancies highlight the need for platelet-focused evaluation methods to set safe design guidelines for cardiovascular nanocarriers. These considerations become particularly relevant for the development of antiplatelet delivery systems based on LPHNPs, where unwanted platelet activation could directly counteract the intended therapeutic effect of the encapsulated payload.

In this work, we employed an integrated *ex vivo* and *in silico* screening strategy to design lipid– polymer hybrid nanoparticles composed of poly(lactic acid) (PLA) and poly(ethylene glycol)-modified lipids —specifically, DSPE-PEG(2000) derivatives with positive (amine-terminated), nominally neutral (methoxy-terminated), or negative (carboxylic acid-terminated) surface charges— for cilostazol (CLZ) delivery. By combining functional assays in washed human platelets and non-equilibrium atomistic steered molecular dynamics (SMD) simulations, we systematically evaluated how nanoparticle surface chemistry influences self-assembly feasibility, nano–platelet interactions, mechanical resistance for drug release, and platelet compatibility. This approach enabled the identification of physicochemical constraints that define safe boundaries at the nano–human platelet interface and revealed a negatively charged LPHNP formulation that exhibits favorable colloidal behavior, preserves platelet compatibility, and is suitable for CLZ delivery.

## 2. MATERIALS AND METHODS

### 2.1. Reagents

Poly (D,L-lactic acid) with a terminal carboxylic acid group (PLA-COOH; inherent viscosity of 0.16–0.25 dL/g) was purchased from Durect Corporation (Pelham, AL, USA). Carboxylic acid-, methoxy-, and amine-terminated derivatives of 1,2-distearoyl-*sn*-glycero-3-phosphoethanolamine-*N*-[(polyethylene glycol)-2000] —commercially designated as DSPE-PEG(2000) carboxylic acid, DSPE-PEG(2000) methoxy, and DSPE-PEG(2000) amine (Cat. Nos. 880135, 880120, and 880128, respectively)— were all obtained from Avanti (Alabaster, AL, USA). Rhodamine B octadecyl ester perchlorate (RhodB) was obtained from Sigma Aldrich (St. Louis, USA). Adenosine 5’-diphosphate (ADP), prostaglandin E1 (PGE-1), theophylline, fibrinogen, and HEPES-Tyrode’s buffer were purchased from Sigma-Aldrich (St. Louis, MO, USA). Cilostazol (CLZ) was obtained from SelleckChem (Houston, TX, USA). Lactate dehydrogenase (LDH) cytotoxicity assay kit was purchased from Cayman Chemical (Ann Arbor, MI, USA). Flow cytometry antibodies against human CD61 (FITC- and PE-conjugated) and PAC-1 (FITC-conjugated) were obtained from BD Pharmingen (BD Biosciences, San Diego, CA, USA).

### 2.2. Synthesis of lipid–polymer hybrid nanoparticles

Lipid–polymer hybrid nanoparticles (LPHNPs) with different surface charges were prepared using a nanoprecipitation/self-assembly method [39,40]. Briefly, PLA-COOH was dissolved in acetonitrile (ACN) at a concentration of 5 mg/mL. Separately, DSPE-PEG(2000) derivatives with different terminal groups (carboxylic acid-, methoxy-, or amine-terminated) were dissolved in a 4% (v/v) aqueous ethanol solution (0.35 mg/mL) and preheated at 65 °C for 3 min. The organic polymer solution was then added dropwise (1 mL/min) to the aqueous lipid solution under gentle stirring at 40 °C. The resulting LPHNPs were maintained under continuous stirring for 4 h at 40 °C to ensure complete evaporation of the organic solvent. The nanoparticles were purified by three successive washing steps (2,000 × g, 10 min each) using Amicon Ultra-15 centrifugal filters with a molecular weight cut-off (MWCO) of 100 kDa and resuspended in ultrapure water to achieve the desired final concentration.

For the preparation of CLZ-loaded LPHNPs, CLZ was first dissolved in dimethyl sulfoxide (DMSO) to obtain a 50 mg/mL stock solution. A 10 µL aliquot of this solution (containing 500 µg of CLZ) was added to 1 mL of PLA-COOH solution (5 mg/mL in ACN), and the resulting mixture was added dropwise to the aqueous lipid solution following the same nanoprecipitation/self-assembly protocol. Fluorescently labeled LPHNPs were prepared similarly by incorporating 3 µL of a Rhodamine B (RhodB) stock solution (10 mg/mL in DMSO) into the organic phase before nanoprecipitation. Finally, the LPHNPs were either stored at 4 °C for immediate use or frozen at −80 °C for subsequent lyophilization.

### 2.3. Physicochemical characterization and stability assays

The hydrodynamic diameter (nm) and zeta potential (ζ, mV) of empty and CLZ-loaded LPHNPs were determined by dynamic light scattering (DLS) and electrophoretic light scattering (ELS), respectively, using a Zetasizer Nano ZS (Malvern Panalytical, Malvern, UK). Samples were prepared by diluting the nanoparticle suspension in 1 mL of ultrapure water and measured at 25 °C.

The morphology of LPHNPs was examined using transmission electron microscopy (TEM) (Hitachi HT7700, Japan). Aliquots (10 µL) of each formulation were deposited onto 300-mesh carbon-coated copper grids and negatively stained with a sterile-filtered aqueous uranyl acetate solution (2% w/v) for 5 min at room temperature. Micrographs were acquired at an accelerating voltage of 80 kV and subsequently analyzed using ImageJ software [41].

To assess colloidal stability, empty and CLZ-loaded LPHNPs were incubated in ultrapure water and maintained at 37 °C under mild agitation for 7 days. The hydrodynamic diameter was monitored at days 0, 1, 2, 3, 4, and 7.

### 2.4. Drug loading and encapsulation efficiency

Drug loading and encapsulation efficiency (EE%) of CLZ were determined using an extraction method adapted from Coimbra *et al*. [42]. Briefly, lyophilized CLZ-loaded formulations (15 mg) were dissolved in 3 mL of ACN. The samples were vortex-mixed, sonicated, and centrifuged, and the resulting supernatants were filtered and analyzed by ultra-performance liquid chromatography (UPLC).

CLZ quantification was performed using an Acquity UPLC system (Waters, Milford, MA, USA) equipped with a binary solvent delivery pump, an autosampler, and a tunable UV detector. Chromatographic separation was achieved using a Waters Acquity BEH C18 column (50 × 2.1 mm, 1.7 μm). The mobile phase consisted of a 60:40 (v/v) acetonitrile:water mixture delivered at a constant flow rate of 0.3 mL/min. The injection volume was 5 μL, with the mobile phase used as the diluent, and the column temperature was maintained at 25 °C. UV detection was performed at 258 nm. For calibration, a standard stock solution was prepared by accurately weighing and dissolving a CLZ reference standard (equivalent to 5 mg of CLZ) in 5 mL of ACN. Working standard solutions were prepared in the concentration range of 4.9–312.5 μg/mL (4.9, 9.8, 19.5, 39.1, 78.1, 156.3, and 312.5 μg/mL).

The encapsulation efficiency (EE%) was calculated as the ratio of experimental to theoretical drug loading, as previously described [43].

### 2.5. *In vitro* drug release

The *in vitro* release profile of CLZ from LPHNPs was evaluated using a Spectra-Por Float-A-Lyzer G2 dialysis system (Sigma-Aldrich, St. Louis, MO, USA). Briefly, 10 mg of CLZ-loaded formulations were introduced into dialysis membrane units with a MWCO of 14 kDa. The dialysis units were sealed and immersed in 5 mL of ultrapure water, placed in an orbital shaker, and maintained at 37 °C. At predetermined time intervals (4, 12, 24, 36, and 48 h), 1 mL of release medium was collected for analysis and immediately replaced with an equal volume of fresh ultrapure water. The concentration of CLZ in the collected samples was quantified by UPLC using the method described above.

### 2.6. Steered molecular dynamics simulations

To probe the relative mechanical resistance to CLZ dissociation across the lipid–polymer interface, non-equilibrium steered molecular dynamics (SMD) simulations were carried out using the Large-scale Atomic/Molecular Massively Parallel Simulator (LAMMPS) [44] and a reactive force field (ReaxFF) [45]. This computational strategy directly follows the methodology validated in our previous study for simulating the self-assembly process of the same LPHNPs used in this work [46]. To obtain a computationally tractable system, a simplified model of the nanoparticle–drug complex was built by replicating the degree of curvature, polymeric core density, and interfacial configurations of the synthesized LPHNPs. The atomistic model comprised ten PLA chains, divided into a rigid and a flexible segment (representing the polymeric chains of the compact core and the region that interacts with the DSPE-PEG lipid chains, respectively). A single CLZ molecule was then placed 3 Å above the center of the core boundary, and two previously minimized and equilibrated DSPE-PEG chains (either methoxy- or carboxylic acid-terminated) [46] were placed above the drug.

The mobile portion of the model (PLA flexible segment, CLZ, and DSPE-PEG chains) was equilibrated for 3 ns at 310 K. Solvent effects were modeled implicitly using Langevin dynamics with a friction coefficient of 30 ps^-1^ and a 1.0 fs timestep [46]. Geometry-dependent partial atomic charges were computed on the fly with the charge equilibration (QEq) method [47], allowing the electrostatic response of the drug to be tracked during conformational changes. Following equilibration, SMD simulations were carried out for each system by pulling the CLZ molecule away from the lipid–polymer region (center-of-mass pulling) along the *z*-axis reaction coordinate. Because the force measured in non-equilibrium SMD depends on the loading rate, a systematic screening of pulling velocities (0.1, 0.5, 1.0, 5.0, and 10.0 Å/ps) and spring constants (50 and 100 kcal/mol·Å^2^) was first carried out for all three systems. Within the 50-ps production window accessible to these reactive simulations, a spring constant of 50 kcal/mol·Å^2^ and a pulling velocity of 5.0 Å/ps allowed complete extraction of the drug while maintaining a smooth, continuous force profile, and were therefore adopted for the final simulations. The maximum pulling force (F_max_) recorded along each trajectory was used as a comparative indicator of interfacial resistance. Since all systems were simulated under identical loading conditions, F_max_ values are interpreted only in relative terms between systems, and not as equilibrium desorption forces, because their absolute magnitude is inherently dependent on the non-equilibrium pulling protocol.

### 2.7. Preparation of washed human platelets

Washed platelet suspensions were obtained from venous blood samples taken from at least fifteen young healthy volunteers. All volunteers signed a written informed consent before the samples were collected. The study protocol was approved by the Ethics Committee of the Universidad de Talca and conducted in accordance with the Declaration of Helsinki. Blood samples were collected by phlebotomy using a tube system containing extraction buffer (PGE-1 5 μg/ml, theophylline 11 μM, pH 7.4). First, the blood samples were centrifuged at 240 × g for 10 min to obtain platelet-rich plasma (PRP). Then, two-thirds of the PRP were collected and centrifuged again for 5 min at 650 × g at 4 °C. The resulting platelet pellet was washed, resuspended in HEPES-Tyrode’s buffer containing PGE-1 (120 nmol/L), and adjusted to 300 × 10^6^ platelets/mL. The platelets were quantified by using the cell counter Bayer Advia 60 Hematology System (Tarrytown, NY, USA). Washed platelets from each volunteer were processed independently in the experiments.

### 2.8. Platelet aggregation assay

Platelet aggregation was determined by light transmission using a lumi-aggregometer (Chrono-Log, Havertown, PA, USA) [48]. Briefly, 480 µL of washed platelets (300 × 10^6^ platelets/mL) were preincubated for 3 min with 20 µL of CLZ (1–50 µM), empty LPHNPs, or CLZ-loaded LPHNPs (8.75, 17.5, and 35.0 µg/mL) in the presence of fibrinogen (275 µg/mL). Platelet aggregation was subsequently induced by the addition of 20 μL of agonist ADP (8 or 4 µM) and recorded for 6 min at 37 °C under constant stirring (1,000 rpm). Maximum aggregation (%) was quantified using AGGRO/LINK software (Chrono-Log).

### 2.9. Platelet disruption assay

Membrane disruption was assessed by measuring LDH release from washed human platelets. The assay was performed according to the manufacturer’s instructions. Briefly, 480 µL of washed platelets (300 × 10^6^ platelets/mL) were incubated with CLZ, empty LPHNPs, or CLZ-loaded LPHNPs for 10 min at 37 °C. Samples were then centrifuged at 1,200 × g for 10 min at 4 °C and the supernatants were collected and incubated with the reaction mixture for 30 min at 37 °C under orbital shaking in a 96-well plate. LDH activity was quantified by measuring formazan dye formation at 490 nm using a Multiskan GO microplate spectrophotometer (Thermo Fisher Scientific, Waltham, MA, USA). Platelet disruption was calculated after normalization to maximum LDH release (positive control) and subtraction of spontaneous LDH release (negative control).

The percentage of platelet disruption was calculated using the following formula:

Platelet disruption (%) = [(Experimental A490 – Negative control A490) / (Positive control A490 – Negative control A490)] x 100,

where Experimental A490 is the absorbance of the washed platelets treated with CLZ or LPHNPs, Negative control A490 corresponds to the vehicle-treated platelets (representing spontaneous LDH release), and Positive control A490 corresponds to completely lysed platelets (representing maximum LDH release).

### 2.10. Flow cytometry study

The activation state of glycoprotein IIb/IIIa (GPIIb/IIIa) on the platelet surface was analyzed by double-label flow cytometry as previously described by Fuentes *et al.* [49]. Briefly, 480 µL of washed platelets (300 × 10^6^ platelets/mL) were incubated with 20 µL of CLZ (1–50 µM), empty LPHNPs, or CLZ-loaded LPHNPs (8.75, 17.5, and 35.0 µg/mL) for 3 min. Samples were then stimulated with ADP (8 or 4 µM) for 6 min at 37 °C. Subsequently, 50 µL aliquots were mixed with saturating concentrations of FITC-conjugated anti-GPIIb/IIIa (PAC-1 clone) and PE-conjugated anti-CD61 antibodies and incubated for 25 min in the dark. Platelet populations were analyzed by acquiring 10,000 events on an Accuri C6 flow cytometer (BD Biosciences, USA). Platelets were identified and gated based on their characteristic light-scattering properties in forward scatter (FSC) vs. side scatter (SSC) plots combined with CD61 positivity.

### 2.11. Nanoparticles–platelet interaction

The association between LPHNPs and washed human platelets was evaluated by flow cytometry. Briefly, 480 µL of washed platelets (300 × 10^6^ platelets/mL) were incubated with 20 µL of RhodB-labeled empty or CLZ-loaded LPHNPs, followed by incubation with saturating concentrations of FITC-conjugated anti-CD61 antibody for 25 min in the dark. Platelet populations were analyzed by acquiring 10,000 events on an Accuri C6 flow cytometer (BD Biosciences, USA). Platelets were identified and gated based on their characteristic light-scattering properties in forward scatter (FSC) vs. side scatter (SSC) plots combined with CD61-FITC positivity.

### 2.12. Statistical analysis

Data are presented as mean ± standard deviation (SD). Three or more independent experiments were performed for each assay. Comparisons between two independent groups were performed using the Mann–Whitney U test, whereas comparisons between two paired groups were analyzed using a two-tailed paired t-test. Differences among multiple paired groups were analyzed using the Friedman test followed by Dunn’s multiple-comparison test. Time-matched controls were included when appropriate. A *p* value < 0.05 was considered statistically significant. Half-maximum inhibitory concentration (IC_50_) values were determined by nonlinear regression analysis. All statistical analyses were conducted using GraphPad Prism 6.0 software (GraphPad Inc., San Diego, CA, USA).

## 3. RESULTS AND DISCUSSION

### 3.1. Formulation screening and self-assembly constraints

The synthesis of the LPHNPs studied here relies on the synchronous nanoprecipitation of the hydrophobic PLA-COOH core alongside the interfacial assembly of amphiphilic DSPE-PEG lipids, a robust methodology widely employed to generate stable core–shell nanostructures [39,40]. To systematically investigate the influence of surface charge on the self-assembly dynamics and colloidal stability of the LPHNPs, we evaluated three distinct terminal lipid charges (positive, amine-terminated; nominally neutral, methoxy-terminated; and negative, carboxyl-terminated). Macroscopic observation revealed that during the self-assembly process, the dropwise addition of the polymeric PLA-COOH solution into the aqueous phase containing the amine-terminated DSPE-PEG—intended to yield the positively charged formulation (NP(+))—resulted in immediate formulation failure, manifested by the development of milky turbidity indicative of rapid aggregation and significant colloidal instability (Fig. S1). Conversely, both neutral and negatively charged formulations (NP(0) and NP(–), respectively) yielded clear, translucent colloidal suspensions. The physicochemical properties of the formulated LPHNPs, including hydrodynamic diameter (or size), polydispersity index (PDI), and zeta potential (ζ-potential), are summarized in Table 1.

**Table 1.**
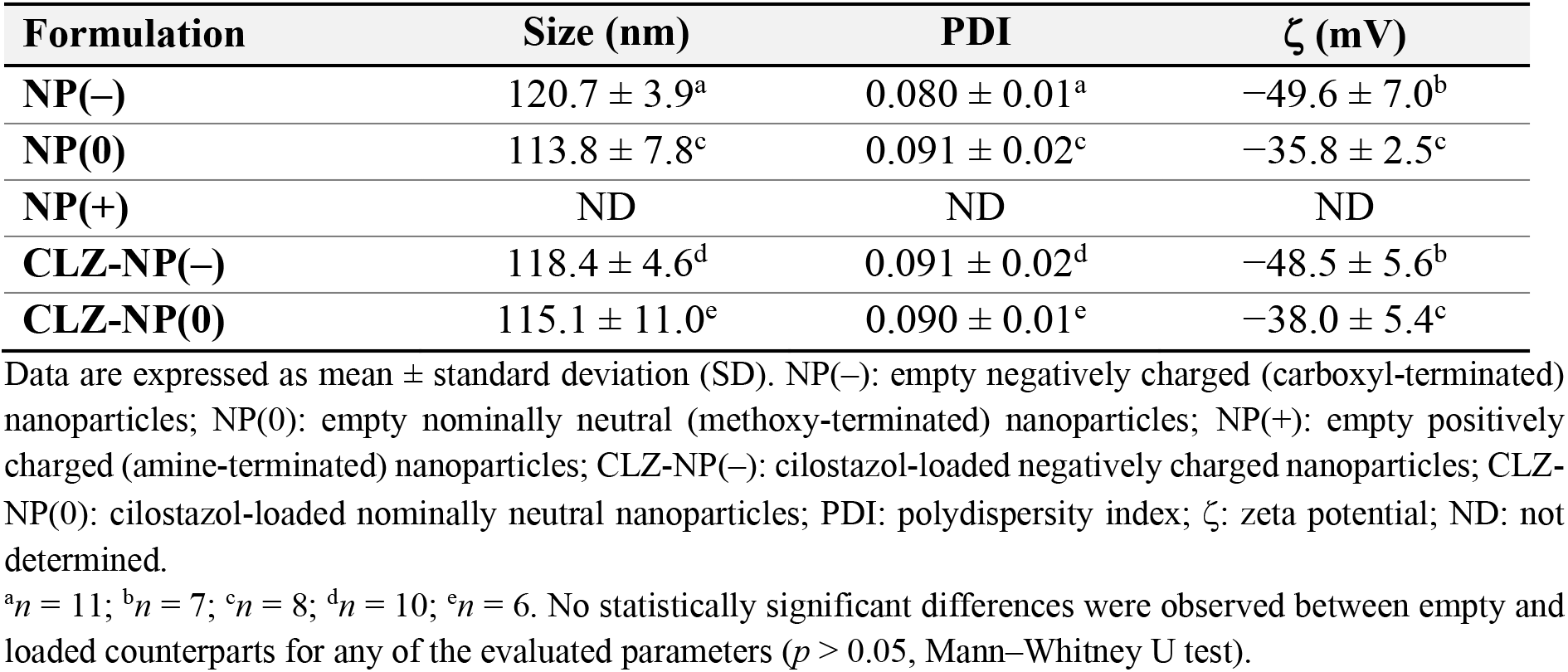
Physicochemical parameters of empty and cilostazol-loaded lipid–polymer hybrid nanoparticles.

The precipitation of the NP(+) formulation can be explained by the interfacial physics involved in the self-assembly process. Under the synthesis conditions, the terminal carboxyl groups of the PLA-COOH polymer core are expected to be predominantly deprotonated (–COO^−^), whereas the primary amine groups in the lipid layer are expected to be predominantly protonated (–NH_3_^+^). The strong electrostatic attraction between these oppositely charged species likely promotes rapid and uncontrolled interfacial complexation. Such interactions may interfere with the cooperative packing of the lipid layer around the polymeric core, thereby compromising colloidal stability and favoring aggregation. Based on this physicochemical limitation, the NP(+) formulation was excluded from subsequent biological evaluations, as it was incompatible with the lipid–polymer hybrid nanostructure investigated in this study.

As summarized in Table 1, the PDI of all neutral and negatively charged formulations (both empty and CLZ-loaded) remained below 0.1, indicating a highly monodisperse particle population. The minor variation in their hydrodynamic diameter upon CLZ loading (∼2 nm) suggests successful drug encapsulation within the polymeric core. TEM micrographs exhibited spherical LPHNPs with uniform sizes and a characteristic heterogeneous distribution of electron density (Fig. 1A–D). The increased electron density observed at the nanoparticle surface is attributed to the DSPE-PEG lipid layer, which absorbs a higher amount of the staining agent. Diameters obtained from TEM analysis showed more pronounced size variations upon CLZ encapsulation compared to those observed by DLS. Specifically, average diameters increased from 58.7 ± 10.0 nm to 89.9 ± 13.9 nm for NP(–) and CLZ-NP(–), respectively, and from 57.4 ± 8.2 nm to 69.6 ± 17.1 nm for NP(0) and CLZ-NP(0), respectively. This discrepancy between DLS and TEM measurements is commonly reported in nanoparticle characterization [50] and reflects the different physical states of the LPHNPs during analysis. DLS measures the nanoparticle diameter in a fully hydrated state [51], where the solvated PEG corona contributes to the measured size and may partially mask structural changes occurring within the polymeric core. In contrast, TEM captures solid-state dimensions under high-vacuum conditions. Under these conditions, the amorphous polymeric core of empty LPHNPs may undergo partial contraction during sample preparation, whereas CLZ incorporation could contribute to preserving a larger apparent particle size by modifying the internal organization or packing density of the polymeric core. Although this interpretation is indirect, the increased TEM diameters observed for CLZ-loaded formulations are consistent with successful drug incorporation within the nanoparticle core. These findings demonstrate that DLS and TEM are complementary techniques, providing valuable insights into colloidal behavior in solution and in the solid state [52].

**Fig. 1.**
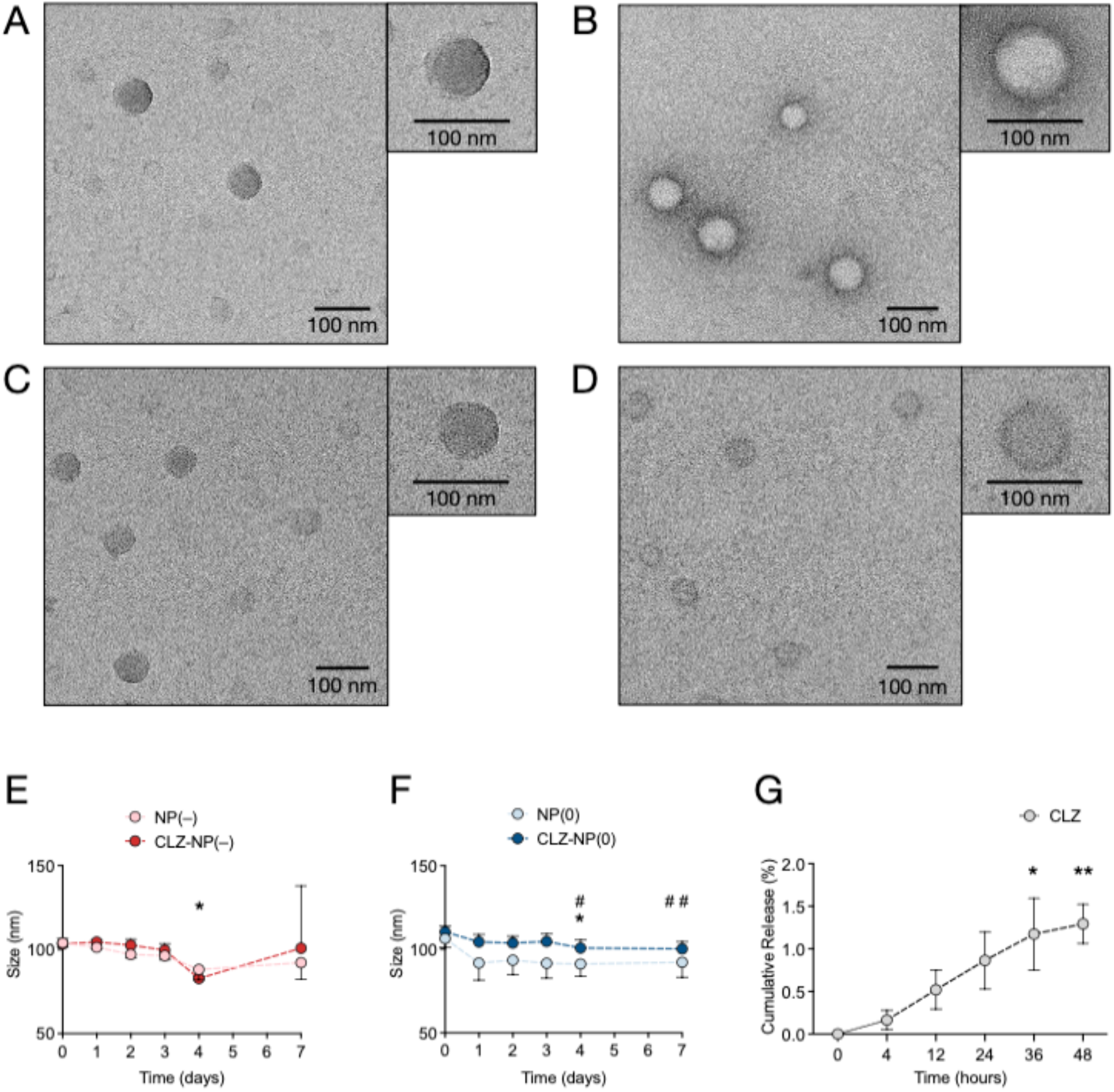
Physicochemical characterization of empty and cilostazol-loaded lipid–polymer hybrid nanoparticles. Representative transmission electron microscopy (TEM) micrographs of (A) empty negatively charged nanoparticles (NP(–)), (B) cilostazol-loaded negatively charged nanoparticles (CLZ-NP(–)), (C) empty neutral nanoparticles (NP(0)), and (D) cilostazol-loaded neutral nanoparticles (CLZ-NP(0)). Insets show higher-magnification images of representative nanoparticles. Scale bars: 100 nm. (E,F) Nanoparticle size stability over 7 days for (E) negatively charged and (F) neutral formulations. (G) *In vitro* cumulative release profile of CLZ from CLZ-NP(0) nanoparticles over 48 hours. In (E), NP(–) exhibited a significant size reduction at day 4 compared with day 0 (*p* = 0.0319). In (F), NP(0) exhibited a significant size reduction at day 4 compared with day 0 (*p* =0.0441), whereas CLZ-NP(0) exhibited significant reductions in size at day 4 (*p* = 0.0441) and day 7 (*p* = 0.0053) relative to day 0. In (G), cumulative CLZ release significantly increased at 36 h (*p* = 0.0441) and 48 h (*p* = 0.0053), whereas the increase observed at 4 h (*p* > 0.9999), 12 h (*p* = 0.9522), or 24 h (*p* = 0.2477) was not statistically significant compared with 0 h. Data are presented as mean ± SD (*n* = 3), and lines connect mean values as a visual guide. Statistical analyses were performed using the Friedman test followed by Dunn’s multiple-comparison test (against day 0 for size stability and against 0 h for cumulative release). ns, not significant; \**p* < 0.05; \*\**p* < 0.01; #*p* < 0.05 and *##*p < 0.01 against day 0 for CLZ-NP(0).

From a biomedical perspective, the average dimensions for both empty and CLZ-loaded formulations suggest a high potential for *in vivo* applications. It has been shown that nanocarriers with diameters of 20–100 nm can circulate in the bloodstream for extended periods and exhibit efficient cellular internalization, facilitating their uptake by various body tissues and organs [53]. In addition, the observed LPHNPs dimensions are optimal for drug delivery, as they are large enough to avoid rapid renal clearance, while the PEGylated surface provides advantages to evade opsonization [54–56].

Although the methoxy-terminated formulations are referred to as NP(0) based on their nominally neutral terminal PEG group, both NP(0) and CLZ-NP(0) exhibited a negative ζ-potential (Table 1). This behavior is likely attributable to the intrinsic negative charges of the phosphate groups present in the DSPE lipid backbone, combined with the deprotonated terminal carboxyl groups of the PLA-COOH core that are not entirely screened by the PEG corona. Similar negative ζ-potential values were previously reported by our group for neutral LPHNPs prepared from the same lipid–polymer components, despite encapsulating a different therapeutic agent, supporting the interpretation proposed here [57]. By contrast, the NP(–) and CLZ-NP(–) formulations exhibited a more pronounced negative ζ-potential (Table 1). This marked shift toward a more negative surface charge directly validates the success of our surface functionalization strategy, confirming the effective and dense exposure of deprotonated carboxylate ions (–COO^−^) at the outermost layer of the hybrid nanostructure. Importantly, the incorporation of CLZ did not significantly alter the surface charge of LPHNPs (Table 1), consistent with the absence of ionizable functional groups in the drug structure. The pronounced negative surface charge across these formulations ensures high colloidal stability and can minimize non-specific binding to cellular membranes via electrostatic repulsion, thereby allowing for a prolonged half-life in circulation [58,59].

### 3.2. Colloidal stability profiles and drug encapsulation

The colloidal stability of neutral and negatively charged LPHNPs was evaluated by monitoring their hydrodynamic diameters over 7 days at 37 °C using DLS. As shown in Fig. 1E and F, a size reduction was observed across all formulations at day 4. For negatively charged nanoparticles, only the empty NP(–) showed a statistically significant decrease, from 103.9 ± 2.4 to 88.1 ± 6.2 nm, while CLZ-NP(–) displayed a similar but non-significant trend (Fig. 1E). In the neutral formulations, both NP(0) and CLZ-NP(0) showed significant size reductions at day 4, with sizes decreasing from 106.4 ± 5.2 to 91.1 ± 7.4 nm and from 110.4 ± 3.5 to 100.7 ± 4.8 nm, respectively. At day 7, only CLZ-NP(0) remained significantly smaller than at day 0, reaching a mean diameter of 100.2 ± 4.2 nm (Fig. 1F). This generalized size reduction by day 4 can be attributed to the progressive thermodynamic relaxation and structural rearrangement of both the polymeric core and the DSPE-PEG layer under thermal incubation. Importantly, the absence of any increase in size over the 7-day study rules out macroaggregation. Since colloidal instability typically drives a pronounced shift toward larger particle sizes, these results confirm that both methoxy- and carboxyl-terminated coatings provide excellent stability under simulated physiological conditions.

The drug loading and encapsulation efficiency (EE%) of the LPHNPs were determined via UPLC after disruption of the polymeric core in the lyophilized formulations. Given that both neutral and negatively charged LPHNPs share an identical hydrophobic PLA-COOH core where the drug is located, the neutral formulation, CLZ-NP(0), was selected as the representative system to establish the drug entrapment and baseline release behavior. The experimental CLZ loading was determined to be 3.7 ± 1.3 % (w/w) relative to a fixed theoretical CLZ loading of 9.09 %, corresponding to an EE% of 40.3 ± 14.3 %. In practical terms, this represents the encapsulation of approximately 37 μg of CLZ per mg of formulation, indicating efficient incorporation of this highly hydrophobic drug into the polymeric core.

*In vitro* drug release profiles obtained by equilibrium dialysis over 48 hours demonstrated progressive, sustained release kinetics. CLZ was released from the LPHNPs at a slow rate, yielding a cumulative release of only 0.5 % during the first 12 hours. After this time window, the release profile transitioned to a linear, zero-order diffusion pattern, reaching approximately 2 % cumulative release at 48 hours (Fig. 1G), corresponding to an absolute drug mass of approximately 7.4 μg in the receptor medium. This controlled release rate is mainly attributed to the high hydrophobicity of CLZ and its strong affinity for the polymeric core. Extrapolation of these *in vitro* findings to physiological environment suggests that *in vivo* release kinetics may follow a more accelerated profile. *In vitro* degradation of the PLA-COOH core is primarily initiated by water infiltration (hydrolytic degradation), which is typically modulated by temperature and pH [60,61], whereas the biological milieu introduces active enzymatic pathways [62]. The presence of circulating esterases in the bloodstream is expected to accelerate degradation of the polymeric core. Therefore, the combined effects of hydrolytic cleavage and enzymatic degradation are likely to result in faster release kinetics compared to those observed *in vitro*.

### 3.3. Interfacial resistance to drug detachment probed by steered molecular dynamics simulations

Non-equilibrium steered molecular dynamics (SMD) simulations were used to compare, at the atomic level, the resistance opposing CLZ detachment from a simplified model of the lipid– polymer interface, as a qualitative complement to the experimental release data (Fig. 2). Under identical loading conditions for all systems, a spring constant (*k* = 50 kcal/mol·Å^2^) and a pulling velocity of 5.0 Å/ps were applied (see Figs. S2–S4 and Methods for details), and the maximum pulling force (F_max_) was determined for three models: (i) an uncoated polymeric core (PLA– CLZ), (ii) a carboxyl-terminated lipid–polymer interface (PLA–CLZ–DSPE-PEG(–), representing the CLZ-loaded negatively charged nanoparticle), and (iii) a methoxy-terminated lipid–polymer interface (PLA–CLZ–DSPE-PEG(0), representing the CLZ-loaded neutral nanoparticle) (Fig. 2A and B).

**Fig. 2.**
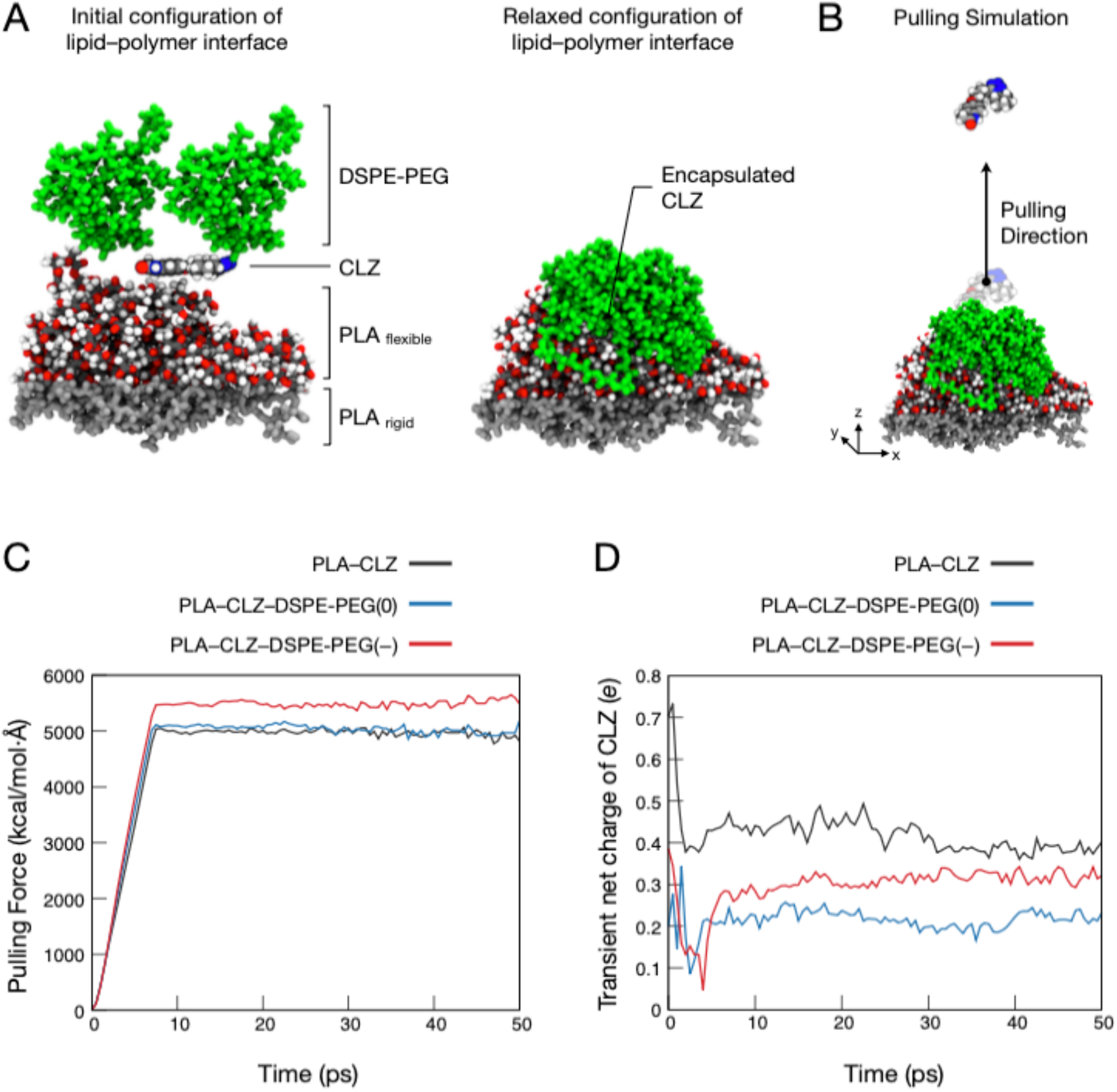
Steered molecular dynamics (SMD) simulations of cilostazol dissociation from lipid– polymer interfaces. (A) Three-dimensional representation of the simplified nanoparticle–drug model before (left) and after a 3-ns equilibration run (right), showing the relaxed lipid–polymer interface and encapsulated cilostazol (CLZ). (B) Schematic representation of the pulling setup used for SMD simulations, indicating the pulling direction applied to CLZ. (C) Pulling force as a function of simulation time during the SMD extraction of CLZ. (D) Time evolution of the transient net charge of CLZ calculated using the charge equilibration (QEq) method. In panels (C) and (D), the black line represents the control system (PLA–CLZ), corresponding to CLZ adsorbed on an uncoated polymeric core; the blue line represents the methoxy-terminated lipid–polymer interface (PLA–CLZ–DSPE-PEG(0)), corresponding to the CLZ-loaded neutral nanoparticle; and the red line represents the carboxyl-terminated lipid–polymer interface (PLA–CLZ–DSPE-PEG(–)), corresponding to the CLZ-loaded negatively charged nanoparticle.

Under these identical non-equilibrium conditions, the control PLA–CLZ system, which lacks the DSPE-PEG lipid layer, exhibited the lowest pulling force (F_max_ = 5,051 kcal/mol·Å) (Fig. 2C). Incorporation of the DSPE-PEG chains increased the force required to extract the drug, with the carboxyl-terminated model (F_max_ = 5,476 kcal/mol·Å) exceeding its methoxy-terminated counterpart (F_max_ = 5,114 kcal/mol·Å) (Fig. 2C). This ordering is consistent with additional steric hindrance and chain entanglement introduced by the lipid layer opposing drug displacement. We note, however, that these are non-equilibrium, protocol-dependent quantities and that the differences between the coated systems are modest; the trends should therefore be regarded as qualitative.

The QEq scheme additionally allowed the electrostatic response of CLZ to be followed during pulling. Although CLZ carries no ionizable groups, its transient net charge fluctuated near zero throughout the trajectories, with a magnitude that depended on the terminal chemistry of the DSPE-PEG lipid layer (Fig. 2D). During the initial pulling phase (0–5 ps), all systems showed sharp fluctuations in the drug’s net charge, coinciding with conformational transitions as the external force perturbed the binding pocket. In the control PLA–CLZ system, the net charge of CLZ shifted toward more positive values, consistent with polarization induced by the electrostatic field of the deprotonated carboxyl groups of the polymeric core. In the two coated models, this excursion was smaller (approximately +0.1 *e* to +0.35 *e*), consistent with partial dielectric screening of the drug by the flexible DSPE-PEG chains. After ∼7.5 ps, coinciding with the maximum pulling forces, the transient charge profiles stabilized (Fig. 2D) as CLZ left the lipophilic core and entered the implicit solvent region. Taken together, these observations suggest that the DSPE-PEG layer may act as both a steric and dielectric barrier to drug displacement, and that the carboxyl-terminated DSPE-PEG coating could provide enhanced sustained CLZ release capacity. While a full thermodynamic quantification of the desorption free energy remains an avenue for future long-timescale studies, the non-equilibrium profiles presented here help elucidate the immediate physical mechanisms driving interfacial drug retention.

### 3.4. *Ex vivo* platelet compatibility

#### 3.4.1. Platelet aggregation profiles and membrane integrity analysis

While CLZ is well known to exert its antiplatelet action by inhibiting platelet aggregation [63], evaluating the potential pro- or anti-thrombotic nature of its nanocarrier is a critical prerequisite for intravascular translation. To establish a baseline pharmacological profile, the antiaggregant effect of free CLZ (1–50 μM) was evaluated using ADP-stimulated washed human platelets. Free CLZ exhibited its classic dose-dependent inhibition of platelet aggregation induced by ADP (8 μM), reducing it from 76.5 ± 3.1 % (control) to 57.4 ± 9.2 % (1 μM), 46.8 ± 8.4 % (5 μM), 33.6 ± 5.2 % (10 μM), and 9.8 ± 2.2 % (50 μM) (Fig. 3A). Expressed as the net percentage of aggregation inhibition, these parameters corresponded to 25.0 ± 12.0 % (1 μM), 38.8 ± 11.0 % (5 μM), 56.1 ± 7.2 % (10 μM), and 87.1 ± 3.0 % (50 μM), yielding an IC_50_ value of 6.8 μM.

**Fig. 3.**
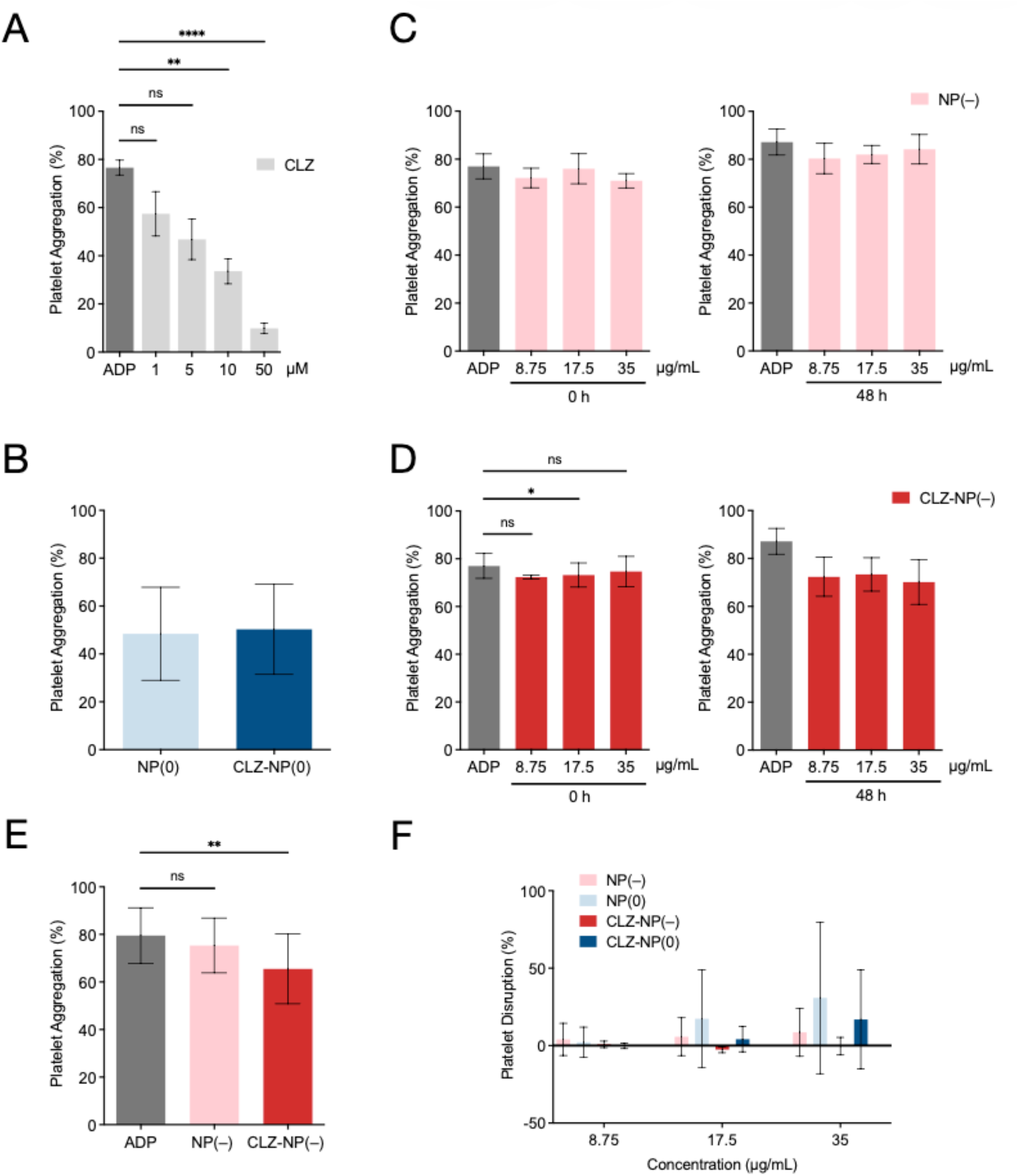
Effect of cilostazol and lipid–polymer hybrid nanoparticles on ADP-induced platelet aggregation and platelet membrane integrity. (A) Effect of cilostazol (CLZ; 1, 5, 10, and 50 µM) on platelet aggregation induced by ADP (8 μM). Data are presented as mean ± SD (*n* = 6). Statistical analysis was performed using the Friedman test followed by Dunn’s multiple-comparison test against ADP. CLZ significantly reduced platelet aggregation at 10 µM (*p* = 0.0041) and 50 µM (*p* < 0.0001), whereas no significant differences were observed at 1 µM (*p* > 0.9999) and 5 µM (*p* = 0.1138) compared with ADP. (B) Platelet aggregation in the presence of empty and CLZ-loaded neutral nanoparticles (NP(0) and CLZ-NP(0), respectively) in the absence of ADP. Data are presented as mean ± SD (*n* = 6). Statistical analysis was performed using a two-tailed paired t-test. No significant difference was detected between NP(0) and CLZ-NP(0) (*p* = 0.5260). (C) Effect of empty negatively charged nanoparticles (NP(–); 8.75, 17.5, and 35 µg/mL) on ADP (8 μM)-induced platelet aggregation without pre-incubation (left) or after 48 h of pre-incubation before platelet exposure (right). Data are presented as mean ± SD (*n* = 3). Statistical analysis was performed using the Friedman test followed by Dunn’s multiple-comparison test against ADP. No significant differences were observed at any NP(–) concentration. (D) Effect of CLZ-loaded negatively charged nanoparticles (CLZ-NP(–); 8.75, 17.5, and 35 µg/mL) on ADP (8 μM)-induced platelet aggregation without pre-incubation (left) or after 48 h of pre-incubation before platelet exposure (right). Data are presented as mean ± SD (*n* = 3). Statistical analysis was performed using the Friedman test followed by Dunn’s multiple-comparison test against ADP. Without pre-incubation (0 h), CLZ-NP(–) significantly reduced platelet aggregation at 17.5 µg/mL (*p* = 0.0342), whereas no significant differences were observed at 8.75 µg/mL (*p* = 0.3415) or 35 µg/mL (*p* = 0.3415). After 48 h of pre-incubation, no significant differences were detected at any concentration (8.75 µg/mL, *p* = 0.1733; 17.5 µg/mL, *p* = 0.3415; 35 µg/mL, *p* = 0.0806). (E) Effect of NP(–) and CLZ-NP(–) formulations (35 μg/mL) on ADP (4 μM)-induced platelet aggregation after 48 h of pre-incubation before platelet exposure. Data are presented as mean ± SD (*n* = 6). Statistical analysis was performed using the Friedman test followed by Dunn’s multiple-comparison test against ADP. CLZ-NP(–) significantly reduced platelet aggregation compared with ADP (*p* = 0.0078), whereas no significant difference was observed for NP(–) (*p* = 0.2978). (F) Assessment of platelet disruption by empty and CLZ-loaded nanoparticles. Platelet disruption was assessed by measuring lactate dehydrogenase (LDH) release from washed human platelets after treatment with NP(–), NP(0), CLZ-NP(–), and CLZ-NP(0) at 8.75, 17.5, and 35 µg/mL for 10 min at 37 °C. Results are expressed as platelet disruption (%), normalized to the vehicle control (negative control, set to 0 %). Data are presented as mean ± SD (*n* = 3). Statistical analysis was performed using the Friedman test followed by Dunn’s multiple-comparison test against the negative control. No statistically significant differences were observed at any nanoparticle concentration. ns, not significant; \**p* < 0.05; \*\**p* < 0.01; \*\*\*\**p* < 0.0001.

Subsequently, the platelet compatibility of the LPHNPs was evaluated. The neutral formulations, NP(0) and CLZ-NP(0), induced spontaneous platelet aggregation in the absence of an external agonist, reaching aggregation levels of 48.4 ± 19.4 % and 50.3 ± 18.8 %, respectively (Fig. 3B). Given this unexpected pro-aggregant effect and its potential thrombotic risk upon systemic administration, the neutral LPHNPs were excluded from all subsequent biological evaluations. In contrast, the negatively charged formulations, NP(–) and CLZ-NP(–), did not induce spontaneous platelet aggregation, highlighting their superior hemocompatibility. Given the restricted release kinetics of CLZ from the nanoparticle core (Fig. 1G), we next investigated whether the encapsulated drug could maintain its antiplatelet activity over time. To this end, both negatively charged formulations were evaluated either immediately after preparation (0 h) or after a 48-h pre-incubation period at 37 °C under constant agitation before platelet exposure, allowing assessment of the functional effect of the released CLZ fraction. Without pre-incubation (0 h), the empty NP(–) formulation did not affect ADP (8 μM)-induced platelet aggregation at any tested concentration (Fig. 3C). Likewise, CLZ-NP(–) did not produce a clear concentration-dependent inhibitory effect, although a statistically significant reduction was observed at 17.5 μg/mL (Fig. 3D). The aggregation values for NP(–) and CLZ-NP(–) were 72.2 ± 4.1 % and 72.3 ± 0.8 % at 8.75 μg/mL; 76.0 ± 6.2 % and 73.2 ± 5.1 % at 17.5 μg/mL; and 71.0 ± 3.0 % and 74.7 ± 6.4 % at 35 μg/mL, respectively.

These observations are consistent with the *in vitro* release profile (Fig. 1G), suggesting that only a limited fraction of CLZ had been released into the medium at the time of platelet exposure. After a 48-h pre-incubation period to allow further drug release, CLZ-NP(–) exhibited a detectable but modest inhibitory effect on platelet aggregation induced by 8 μM ADP (Fig. 3D), yielding aggregation values of 72.3 ± 8.1 %, 73.3 ± 7.0 %, and 70.2 ± 9.4 % at 8.75, 17.5, and 35 μg/mL, respectively, compared with the stimulated control (87.2 ± 5.4 %). Considering that this limited response is likely associated with the slow-release kinetics of CLZ rather than insufficient drug loading, we hypothesized that the antiplatelet activity of the released fraction could be partially masked by the relatively high ADP concentration (8 μM) used in these experiments. To test this hypothesis, the 48-h pre-incubated negatively charged formulations were subsequently evaluated against a lower ADP stimulus (4 μM) at the maximum nanoparticle concentration tested (35 μg/mL). Under these conditions, CLZ-NP(–) significantly reduced platelet aggregation to 65.5 ± 14.7 % compared with 79.5 ± 11.7 % in the ADP-stimulated control (Fig. 3E). In contrast, the empty formulation, NP(–), produced no significant change in the aggregation profile (75.3 ± 11.5 %), indicating that the observed antiplatelet effect was attributable to the released CLZ fraction.

To determine whether the spontaneous platelet aggregation induced by the neutral LPHNPs was associated with a loss of platelet membrane integrity, lactate dehydrogenase (LDH) leakage assays were performed using the same exposure times employed in the platelet aggregation experiments. Previous studies have shown that nanocarriers of the same size but with different surface charges can interact with cell membranes through distinct mechanisms, in some cases causing local membrane perturbation upon contact [32,64,65]. In platelets, these interactions are particularly complex due to the large number of membrane receptors and intracellular signaling pathways involved in activation processes [66]. Membrane disruption or nanoparticle-induced pore formation may promote the release of intracellular granule contents, including pro-thrombotic mediators, thereby contributing to unintended platelet activation and aggregation [67].

The LDH levels detected in the supernatants of washed human platelets treated with empty and CLZ-loaded LPHNPs at concentrations of 8.75, 17.5, and 35 μg/mL showed no statistically significant differences compared with the negative control (Fig. 3F). Specifically, the negatively charged formulations, NP(–) and CLZ-NP(–), exhibited minimal platelet disruption values of 3.9 ± 10.3 % and 0.8 ± 2.1 % at 8.75 μg/mL; 5.8 ± 12.2 % and −2.6 ± 1.9 % at 17.5 μg/mL; and 8.6 ± 15.4 % and −0.3 ± 5.6 % at 35 μg/mL, respectively. Similarly, the neutral formulations, NP(0) and CLZ-NP(0), showed platelet disruption values of 2.2 ± 9.7 % and −0.1 ± 1.8 % at 8.75 μg/mL; 17.3 ± 31.6 % and 4.2 ± 8.1 % at 17.5 μg/mL; and 30.8 ± 48.9 % and 16.9 ± 31.9 % at 35 μg/mL, respectively (Fig. 3F). Although a numerical increase in LDH release was observed for NP(0) at the highest concentration tested, this trend did not reach statistical significance because of the high variability among samples. Likewise, free CLZ (1– 50 μM) did not induce significant LDH release at any tested concentration, indicating that the drug itself does not compromise platelet membrane integrity under the experimental conditions evaluated (Fig. S5).

Collectively, these findings suggest that the spontaneous platelet aggregation induced by neutral LPHNPs is unlikely to be mediated by direct structural damage or membrane disruption. Instead, the methoxy-functionalized surface of NP(0) and CLZ-NP(0) may promote platelet activation through receptor-mediated or other non-lytic mechanisms, consistent with previous observations reported for multiwalled carbon nanotubes [36]. Such interactions could stimulate intracellular signaling pathways that ultimately lead to granule secretion and platelet aggregation.

#### 3.4.2. Platelet activation and nanoparticle–platelet association

Given that platelet aggregation is preceded by early activation events, we next investigated whether negatively charged LPHNPs modulate the activation state of the glycoprotein IIb/IIIa (GPIIb/IIIa) integrin complex using flow cytometry. GPIIb/IIIa was selected as the primary activation marker because it integrates multiple signaling pathways that culminate in aggregation [4,68,69], thereby serving as an effective marker for assessing nanoparticle interactions with the platelet membrane.

To establish the baseline pharmacological behavior on this marker, washed human platelets were first exposed to free CLZ (1–50 μM). The free drug showed a modest, non-linear inhibitory effect on ADP (8 μM)-induced GPIIb/IIIa activation, reducing expression levels from 81.6 ± 9.3 % in the stimulated control to 67.3 ± 17.5 % at 1 μM and 58.3 ± 22.9 % at 5 μM. Interestingly, this inhibitory effect was not dose-dependent. A statistically significant reduction relative to the stimulated control was observed only at 10 μM (49.9 ± 24.8 %), whereas treatment with 50 μM produced a plateau-like response with increased data dispersion (58.5 ± 30.4 %) (Fig. 4A). This limited efficacy on the initial conformational change of GPIIb/IIIa is consistent with the well-characterized pharmacodynamics of CLZ. As previously demonstrated by Inoue *et al*. [70] and further supported by Ito *et al.* [71], although cAMP-elevating agents like CLZ strongly inhibit alpha-granule exocytosis, their ability to suppress the early inside-out signaling required for GPIIb/IIIa activation is comparatively weaker, particularly under direct purinergic stimulation with ADP.

**Fig. 4.**
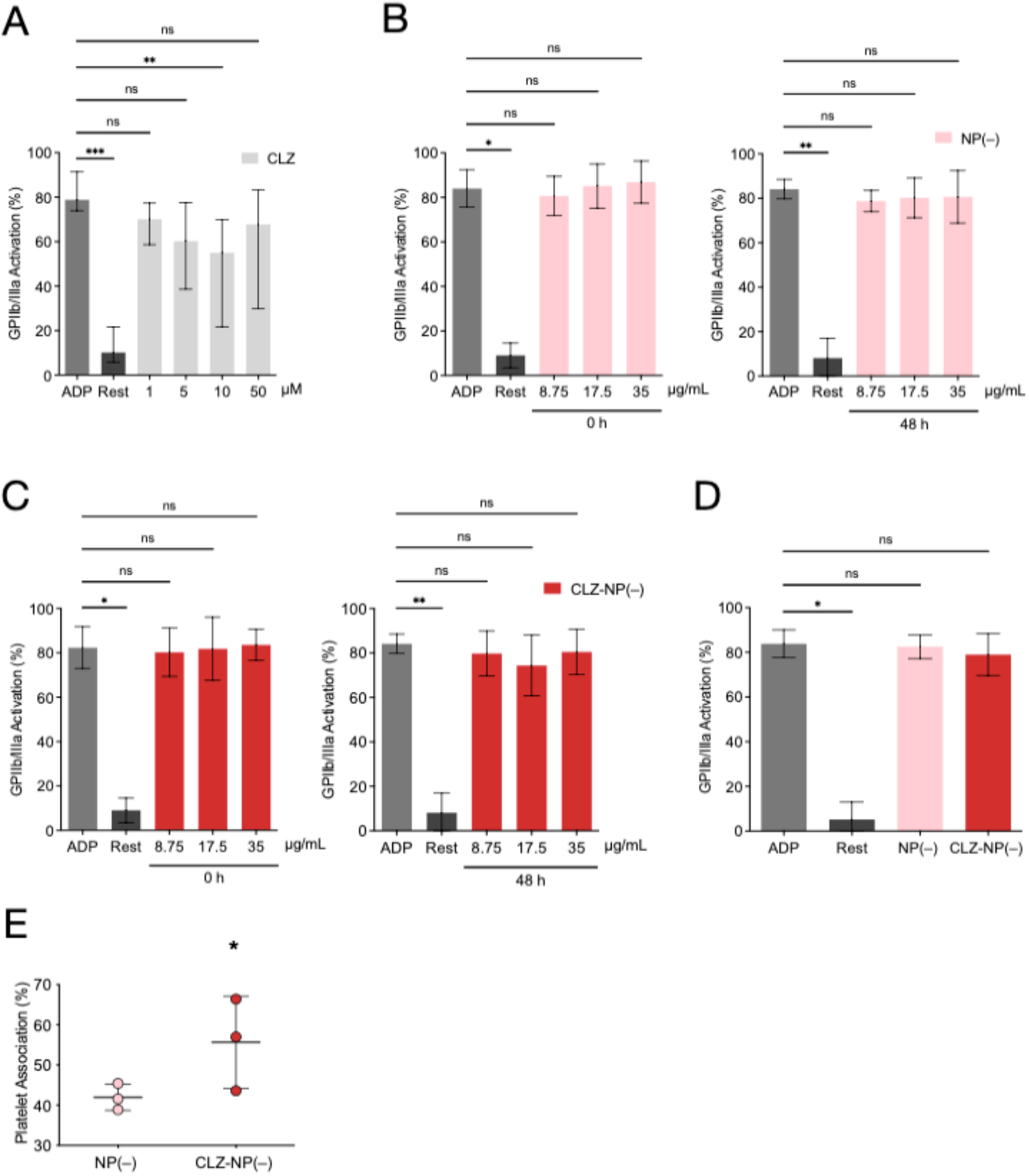
Effect of cilostazol and lipid–polymer hybrid nanoparticles on ADP-induced platelet activation (GPIIb/IIIa activation) and evaluation of nanoparticle–platelet association. (A) Effect of cilostazol (CLZ; 1, 5, 10, and 50 µM) on GPIIb/IIIa activation induced by ADP (8 μM). Data are presented as mean ± SD (*n* = 6). Statistical analysis was performed using the Friedman test followed by Dunn’s multiple-comparison test against ADP. Resting platelets (Rest) exhibited significantly lower GPIIb/IIIa activation than ADP-stimulated platelets (*p* = 0.0002). CLZ significantly reduced GPIIb/IIIa activation at 10 µM (*p* = 0.0060), whereas no significant differences were observed at 1 µM (*p* = 0.6141), 5 µM (*p* = 0.1538), or 50 µM (*p* = 0.3204) compared with ADP. (B) Effect of empty negatively charged nanoparticles (NP(–); 8.75, 17.5, and 35 µg/mL) on ADP (8 μM)-induced GPIIb/IIIa activation without pre-incubation (left) or after 48 h of pre-incubation before platelet exposure (right). Data are presented as mean ± SD (*n* = 6). Statistical analysis was performed using the Friedman test followed by Dunn’s multiple-comparison test against ADP. Resting platelets (Rest) exhibited significantly lower GPIIb/IIIa activation than ADP-stimulated platelets (0 h, *p* = 0.0423; 48 h, *p* = 0.0041), whereas no significant differences were observed at any NP(–) concentration. (C) Effect of CLZ-loaded negatively charged nanoparticles (CLZ-NP(–); 8.75, 17.5, and 35 µg/mL) on ADP (8 μM)-induced GPIIb/IIIa activation without pre-incubation (left) or after 48 h of pre-incubation before platelet exposure (right). Data are presented as mean ± SD (*n* = 6). Statistical analysis was performed using the Friedman test followed by Dunn’s multiple-comparison test against ADP. Resting platelets (Rest) exhibited significantly lower GPIIb/IIIa activation than ADP-stimulated platelets (0 h, *p* = 0.0247; 48 h, *p* = 0.0041), whereas no significant differences were observed at any CLZ-NP(–) concentration. (D) Effect of NP(–) and CLZ-NP(–) formulations (35 μg/mL) on ADP (4 μM)-induced GPIIb/IIIa activation after 48 h of pre-incubation before platelet exposure. Data are presented as mean ± SD (*n* = 4). Statistical analysis was performed using the Friedman test followed by Dunn’s multiple-comparison test against ADP. Resting platelets (Rest) exhibited significantly lower GPIIb/IIIa activation than ADP-stimulated platelets (*p* = 0.0185), whereas no significant differences were observed for NP(–) (*p* > 0.9999) or CLZ-NP(–) (*p* = 0.8200). (E) Association between platelet membranes and fluorescently labeled NP(–) and CLZ-NP(–) formulations. Fluorescent-positive platelets were quantified and expressed as platelet association (%), normalized to the vehicle control (negative control, set to 0 %). Data are presented as mean ± SD (*n* = 3), with individual data points overlaid. Statistical analysis was performed using the Friedman test followed by Dunn’s multiple-comparison test against the negative control. CLZ-NP(–) exhibited significantly increased platelet association relative to the negative control (*p* = 0.0286), whereas no significant difference was observed for NP(–) (*p* = 0.4413). ns, not significant; \**p* < 0.05; \*\**p* < 0.01; \*\*\**p* < 0.001.

Following the experimental protocol established for the platelet aggregation assays, negatively charged LPHNPs were evaluated either immediately (0 h) or after a 48-h pre-incubation period at 37 °C under constant agitation to allow progressive CLZ release before platelet exposure. Across all tested concentrations (8.75, 17.5, and 35 μg/mL), neither the empty NP(–) nor the drug-loaded CLZ-NP(–) formulations significantly altered ADP (8 μM)-induced GPIIb/IIIa activation, regardless of the pre-incubation period (Fig. 4B and C). Without pre-incubation (0 h), platelet activation levels remained comparable to those of the ADP-stimulated control, ranging from 81.8 % to 86.9 % versus 80.7 ± 10.4 %. After 48 h of pre-incubation, a modest reduction in activation levels was observed (74.4 % – 80.6 %), although these values did not differ significantly from the stimulated control (84.1 ± 4.4 %). In addition, under a lower ADP stimulus (4 μM) and using the highest nanoparticle concentration tested (35 μg/mL after 48 h of pre-incubation), neither formulation significantly reduced GPIIb/IIIa activation, yielding values of 82.6 ± 5.3 % for NP(–) and 79.0 ± 9.3 % for CLZ-NP(–) compared with 83.8 ± 6.2 % in the stimulated control (Fig. 4D). Therefore, these findings suggest that the fraction of CLZ released from CLZ-NP(–) is sufficient to attenuate platelet aggregation but insufficient to markedly impair the early inside-out signaling events required for GPIIb/IIIa activation.

Finally, the association between rhodamine B (RhodB)-labeled negatively charged LPHNPs and platelets was evaluated by flow cytometry. As shown in Fig. 4E, exposure to NP(–) and CLZ-NP(–) resulted in platelet association values of 41.9 ± 3.3 % and 55.7 ± 11.5 %, respectively. Only CLZ-NP(–) exhibited a statistically significant increase compared with the negative control. Despite this efficient nanoparticle–platelet association, neither formulation induced membrane damage or spontaneous platelet activation under the experimental conditions evaluated. Collectively, these findings suggest that negatively charged LPHNPs establish a biocompatible interface with platelet membranes, allowing efficient nanoparticle association while preserving platelet quiescence and membrane integrity.

## 4. CONCLUSIONS

This study demonstrates that the surface chemistry of lipid–polymer hybrid nanoparticles based on PLA and DSPE-PEG is a critical factor for their structural stability, drug retention properties, and human platelet compatibility. The nanoprecipitation/self-assembly method successfully generated highly monodisperse neutral and negatively charged formulations, whereas positively charged functionalization caused immediate macroscopic collapse due to chaotic electrostatic complexation during assembly. This finding highlights the importance of interfacial electrostatic balance during the formation of hybrid nanostructures.

Neutral nanoparticles achieved colloidal stability but caused harmful, spontaneous platelet aggregation through non-lytic membrane activation. In contrast, negatively charged nanoparticles emerged as the only viable, hemocompatible carriers, as they maintained excellent colloidal stability, preserved platelet membrane integrity, and exhibited platelet anti-aggregant effects via sustained release of encapsulated cilostazol. Non-equilibrium steered molecular dynamics simulations further suggest that surface functionalization modulates the interfacial resistance to drug extraction, with the negatively charged lipid coating showing the highest resistance under identical pulling conditions.

Finally, these findings highlight the need for rigorous platelet compatibility testing using experimental–computational approaches for any nanomaterial intended for blood-contacting applications and establish negatively charged lipid–polymer hybrid nanoparticles as a promising platform for controlled antithrombotic therapy in the bloodstream.

## ACKNOWLEDGMENTS

M.F.M. acknowledges support from CONICYT-PFCHA/Doctorado Nacional/2014-21140225. C.V. gratefully acknowledges financial support from the Chilean National Agency for Research and Development (ANID) through FONDECYT Regular Grant No. 1241371 and from the Center for Nanoscience and Nanotechnology (CEDENNA) under award CIA250002. B.A.C. acknowledges financial support from ANID through FONDECYT Postdoctoral Grant No. 3170938. M.M.M. acknowledges support from FONCyT PICT-A-2020–2943, CONICET PIP 1122020010, and the PAGE Program from SECyT, Universidad Nacional de Córdoba. J.S.M. acknowledges support from ANID Postdoctoral Grant No. 3240414. E.F. acknowledges support from FONDECYT Initiation Grant No. 11140142. I.P. acknowledges support from the PIEI-ES Program at the Universidad de Talca. E.F. and I.P. also acknowledge the financial support provided by ANID/FONDECYT Regular Grant No. 1260612 and ANID/FONDECYT Regular Grant No. 1260773. The authors also thank the Interuniversity Center for Healthy Aging (Code RED211993) and the Red Interuniversitaria de Envejecimiento Saludable, Latinoamérica y Caribe (RIES-LAC) for their support. Computational resources were provided by the Centro de Cómputo de Alto Desempeño (CCAD-UNC), Universidad Nacional de Córdoba (ccad.unc.edu.ar) and Escuela de Ingeniería Civil en Bioinformática, Universidad de Talca.

