## Supplementary material for "Surface charge tuning of lipid–polymer hybrid nanoparticles for optimized cilostazol delivery and platelet compatibility": SI material

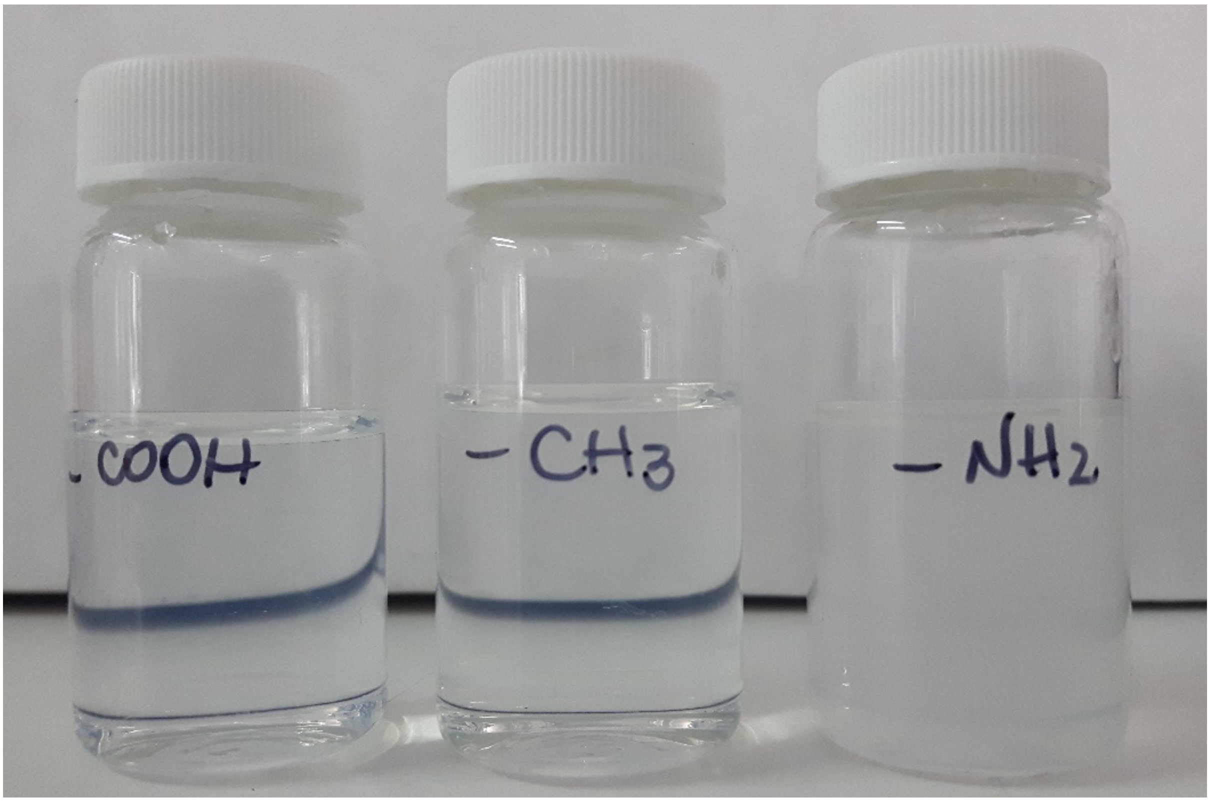


**Figure S1. Visual appearance of surface-functionalized lipid–polymer hybrid nanoparticles (LPHNPs) at room temperature.** Vials show aqueous dispersions of carboxyl-terminated (negatively charged), methoxy-terminated (neutral), and amine-terminated (positively charged) formulations. A distinct milky turbidity is observed only in the amine-terminated formulation.

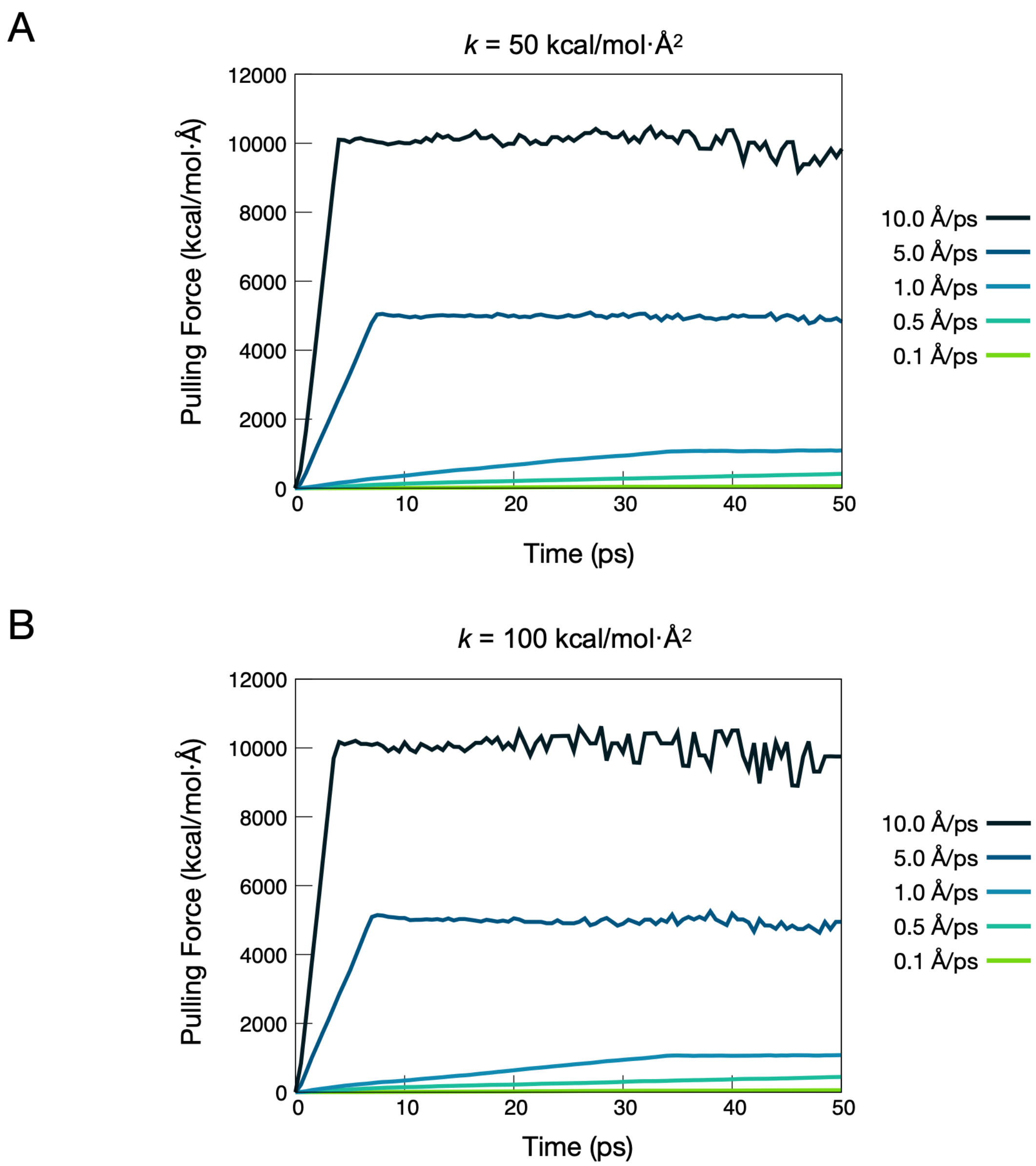


**Figure S2. Systematic parameter screening for Steered Molecular Dynamics (SMD) simulations of the PLA–CLZ model.** Pulling force profiles as a function of simulation time evaluated across varying pulling velocities (0.1, 0.5, 1.0, 5.0, and 10.0 Å/ps) using a harmonic spring constant of either (A) $k$ = 50 kcal/mol·Å$^2$ or (B) $k$ = 100 kcal/mol·Å$^2$.

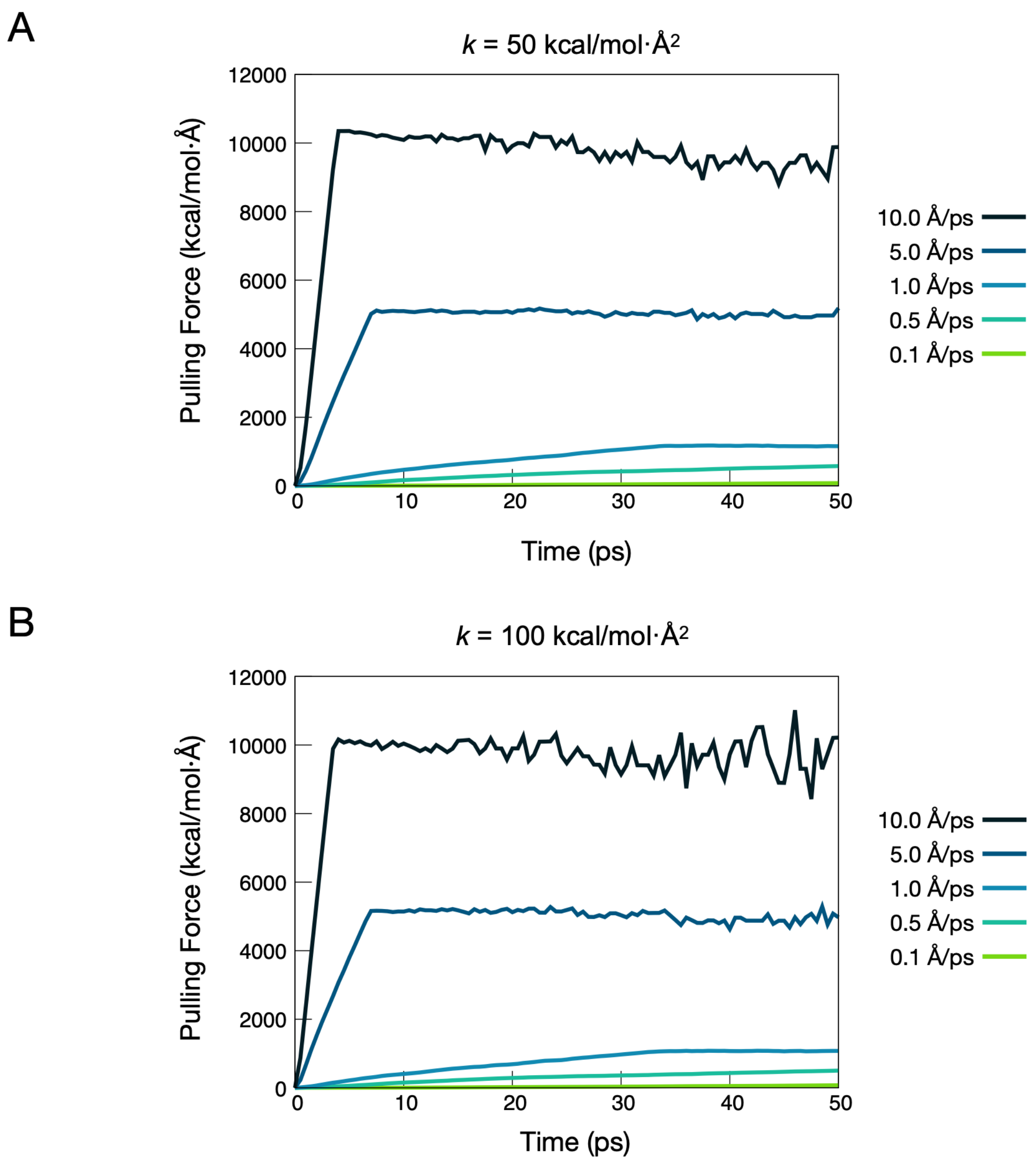


**Figure S3. Systematic parameter screening for Steered Molecular Dynamics (SMD) simulations of the PLA–CLZ–DSPE-PEG(0) model.** Pulling force profiles as a function of simulation time evaluated across varying pulling velocities (0.1, 0.5, 1.0, 5.0, and 10.0 Å/ps) using a harmonic spring constant of either (A) $k$ = 50 kcal/mol·Å$^2$ or (B) $k$ = 100 kcal/mol·Å$^2$.

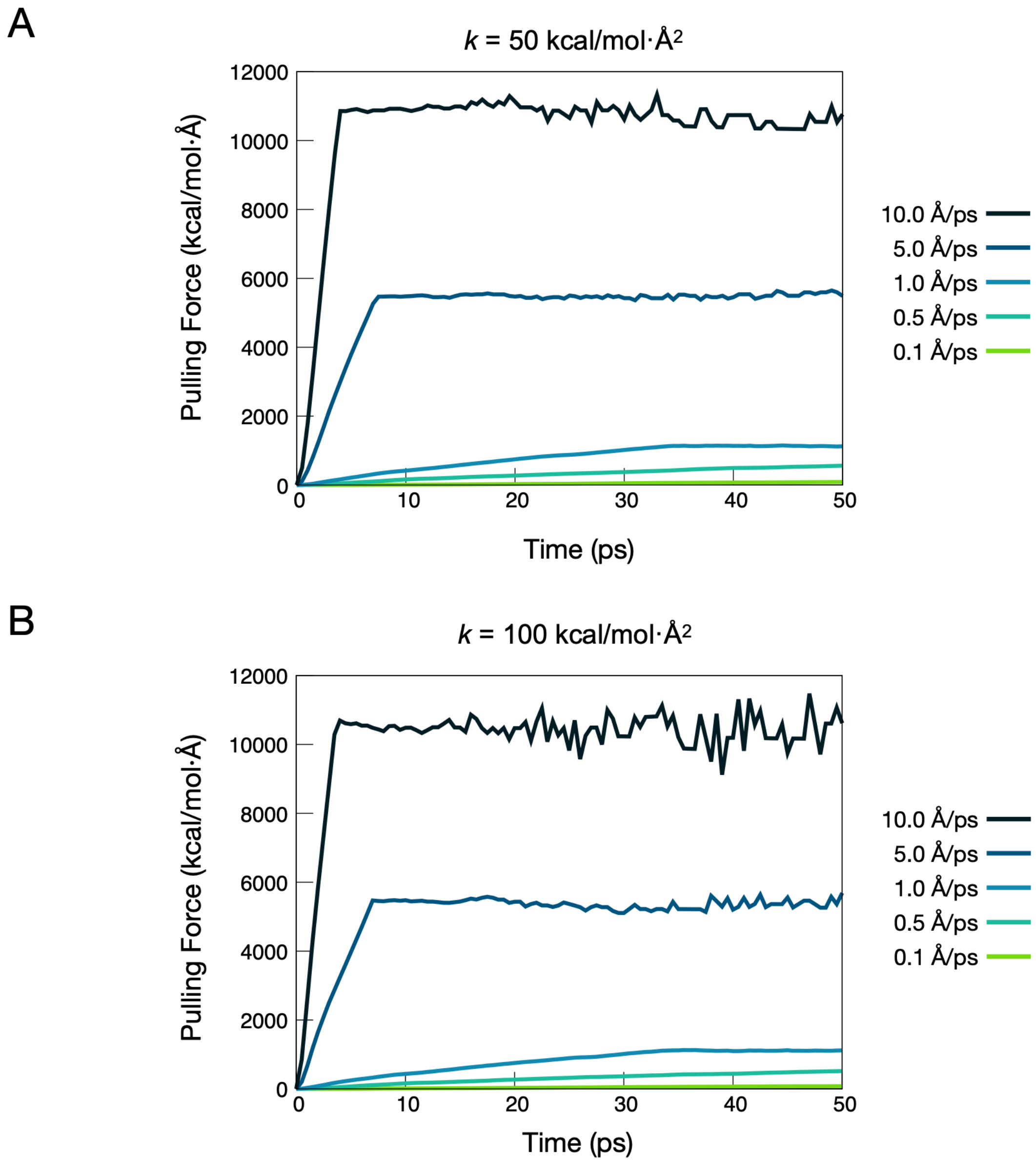


**Figure S4. Systematic parameter screening for Steered Molecular Dynamics (SMD) simulations of the PLA–CLZ–DSPE-PEG(–) model.** Pulling force profiles as a function of simulation time evaluated across varying pulling velocities (0.1, 0.5, 1.0, 5.0, and 10.0 Å/ps) using a harmonic spring constant of either (A) $k$ = 50 kcal/mol·Å$^2$ or (B) $k$ = 100 kcal/mol·Å$^2$.

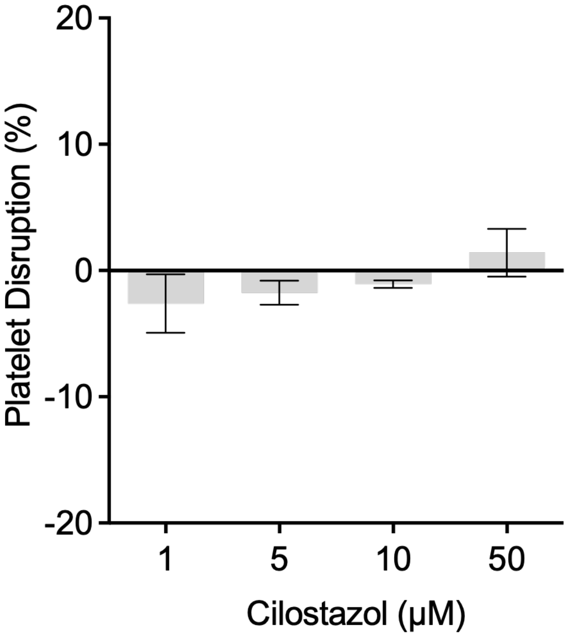


**Figure S5. Assessment of platelet disruption by cilostazol.** Platelet disruption was assessed by measuring lactate dehydrogenase (LDH) release from washed human platelets after treatment with cilostazol (1, 5, 10, and 50 μM) for 10 min at 37 °C. Results are expressed as platelet disruption (%), normalized to the vehicle control (negative control, set to 0 %). Data are presented as mean ± SD ($n$ = 3). Statistical analysis was performed using the Friedman test followed by Dunn's multiple-comparison test against the negative control. No statistically significant differences were observed at any cilostazol concentration.
